# Computational modelling of specific serotonin reuptake inhibitor inhibition on the TREK-2 channel

**DOI:** 10.1101/2024.05.01.592054

**Authors:** Narawit Pratueangsukpong, Tanadet Pipatpolkai

## Abstract

The TREK-2 is a mechanosensitive potassium channel in the two-pore (K2P) potassium channel subfamily. Recent studies of the TREK-2 channel with norfluoxetine reveal that norfluoxetine stabilises a conformation with a lower open probability and disrupts channel gating through a selectivity filter. In addition, multiple specific serotonin reuptake inhibitors (SSRIs) have previously been shown to inhibit the TREK channels subfamily. However, the mechanism of lipid-like SSRI inhibition to the TREK-2 channel is currently unclear. Using molecular dynamic simulation, we show that fluoxetine and escitalopram share the same binding pocket on the TREK-2 channel. We show that fluoxetine inhibits the TREK-2 channel using POPC lipid and directly disrupts the channel gating at the selectivity filter, while escitalopram is a traditional pore blocker, which also disrupts the selectivity filter gating but without POPC dependent inhibition. In addition, we show that both fluoxetine and escitalopram prevent a down-to-up transition when the pressure is applied to the system, showing a conserved mechanism of TREK-2 inhibition. Together, our work reveals mechanistic insight into TREK-2 channel inhibition by lipid-like antidepressants, which could further shed light on rational drug design in the future.

## Introduction

The TREK subfamily of the two-pore (K2P) potassium channel are mechanosensitive potassium channels and are responsible for maintaining cellular resting potentials and other excitable activities within the central and peripheral nervous system (Niemeyer et al. 2016). TREK channels are gated by pressure, chemical stimuli, and lipid environment, which influence their potassium ion conductance (Djillani, Mazella, et al. 2019). Due to their diverse expression natures, they are one of the more prominent pharmacological targets for anaesthesia, chronic pain and depression (Noël, Sandoz, and Lesage 2011). Multiple antidepressants, such as fluoxetine, R-citalopram, escitalopram, and amitriptyline, which are specific serotonin reuptake inhibitors (SSRI), have also been shown to inhibit the TREK channel superfamily (Kennard et al. 2005; Al-Moubarak, Veale, and Mathie 2013; Lin et al. 2015; Park et al. 2016). Understanding the mechanisms of antidepressant inhibitions on the TREK channel will aid further therapeutic design in the future (Djillani, Pietri, et al. 2019).

Recent structures of TREK channels (both TREK-1 and TREK-2) and electrophysiological studies have elucidated the gating mechanisms of the TREK-2 channel (Dong et al. 2015; Lolicato et al. 2017b; Schmidpeter et al. 2023). Until now, two distinct conformations, up and down, of the TREK channel subfamily have been solved. These two distinct conformational changes are commonly observed amongst the K2P channel superfamily (Schreiber, Düfer, and Seebohm 2022). These two conformations are defined based on the position of the TM4 helix, where the up conformation (from TREK-1) has the M4 helix more parallel towards the membrane. Electrophysiological studies show that both conformations of K2P channels are conductive and structurally; neither conformation has a clear bundle crossing gate for hydrophobic gating (Brohawn, del Mármol, and MacKinnon 2012; Miller and Long 2012; Lolicato et al. 2014). Similar to the previously C-type inactivation observed in voltage-gated potassium channels, the TREK channel subfamily is gated at the selectivity filter (Zilberberg, Ilan, and Goldstein 2001; Piechotta et al. 2011; Nematian-Ardestani et al. 2020). Electrophysiological and computational studies have highlighted that the down conformation shows a lower open probability than the up conformation due to the gating event selectivity filter (Schewe et al. 2016). This suggested a strong coupling between the conformation of the M4 helix, and the dynamics of the selectivity filter. In addition, distortion of the selectivity filter has also been shown to be associated with the deeper-down conformation, which is shown to be non-conductive (Zhang et al. 2022). However, this specific conformation of the TREK channel has not been solved structurally. All-atom molecular dynamic simulations have demonstrated that the down-to-up conformational change occurs when negative pressure is applied to the system (Aryal et al. 2017). Thus, by applying the pressure to the system, the channel increases its open probability.

Pharmacological targeting of the TREK channel has been heavily studied structurally, electrophysiologically and computationally. SSRI antidepressants, including fluoxetine, escitalopram and amitriptyline, have been previously shown to inhibit the TREK-2 channel with high efficacy (Kennard et al. 2005; Park et al. 2016). Structural studies have highlighted the binding site of norfluoxetine, a fluoxetine metabolite, at the lipid fenestration gate of the TREK2 channel (Dong et al. 2015; Lolicato et al. 2017a). This site is only available for drug binding when the channel is in the down conformation, suggesting the state-dependent TREK channel inhibition. The sequence at the lipid fenestration site is highly conserved amongst all the TREK channels (Schmidpeter et al. 2023). Computational studies have highlighted that binding of the fluoxetine at the binding site has prevented the down-to-up conformational change when the pressure is applied to the channel (Clausen et al. 2017). Single-channel recording has also suggested some mechanistic insights, highlighting that the blocking of the channel by norfluoxetine is not dependent on its charge, but the drug influences the selectivity filter gating (Proks et al. 2021).

In this study, we used molecular dynamics simulations to elucidate the mechanism of fluoxetine and escitalopram inhibition on the TREK-2 channel. Our simulations show that fluoxetine and escitalopram share the same binding pocket, one at the fenestration site and one below the selectivity filter - blocking the channel like a traditional blocker. As fluoxetine and escitalopram bind to the channel, the selectivity filter is distorted and, thus, likely to affect the gating of the TREK-2 channel. In addition, we also show that fluoxetine introduces lipid-dependent blocking of the channel at the lower gate of the TREK-2 channel. Last, we show that both fluoxetine and escitalopram prevent channel down-to-up conformational transition. Together, this work may suggested an overall inhibitory mechanism for lipid-like SSRI on TREK channels subfamily.

## Results

### Fluoxetine and escitalopram move towards the central cavity of the pore

Previous studies have shown that Fluoxetine (FLX) bind to the lipid fenestration site of the down-state TREK2 channel and, thus, has the potential to behave as the channel pore blocker (Lolicato et al. 2017a). We are interested in the behaviour of lipophilic SSRIs within the lipid fenestration site. First, we conducted 500 ns molecular dynamics simulations of the TREK2 channel with either two fluoxetine (n = 4) or two escitalopram (n = 8) molecules bound in the fenestration site, similar to the crystallographic pose of norfluoxetine (Figure 1A). As TREK-2 is a symmetrical dimer and our simulation is too short to display any co-operativity between the two subunits, we expect the results from each subunit to be independent. For both fluoxetine and escitalopram, we showed that after 500 ns of simulation, one of the molecules moves to the pore’s central cavity, whilst the other molecule remains bound in the binding site. We then conducted the contact analysis to determine whether the amino acid contacting residues change as drugs move from their initial binding site. Interestingly, both drugs show similar contacting residues, highlighting their similar nature within the binding pocket (Figure 1B). In this study, we define contacting residue as amino acids within 4 Å of the drug molecules more than 60% of the simulation time. Despite its dynamical motion towards the pore’s central cavity, we show that the contacting residues remain unchanged. In fluoxetine, we show that F316 is making contact with the trifluoromethyl group, and the amine group interacts with T172 and T281 via hydrogen bonding (Figure 1B,C). The rest of the contacting residues are the nearby hydrophobic amino acids (I197, F200, L204, L279, T280, L313, A317 and L320) which stabilise the aromatic ring within the fluoxetine molecule. In escitalopram, T172 interacts with fluorine, while F316 interacts with the nitrile group. The rest of the amino acids interact with the molecules’ hydrophobic region. This highlights the conserved feature of the two drugs, i.e. the hydrophobic ring region and the hydrogen bond acceptor and their interaction in the binding pocket. This suggested that fluoxetine and escitalopram binding pocket is shared between them.

**Figure 1.**
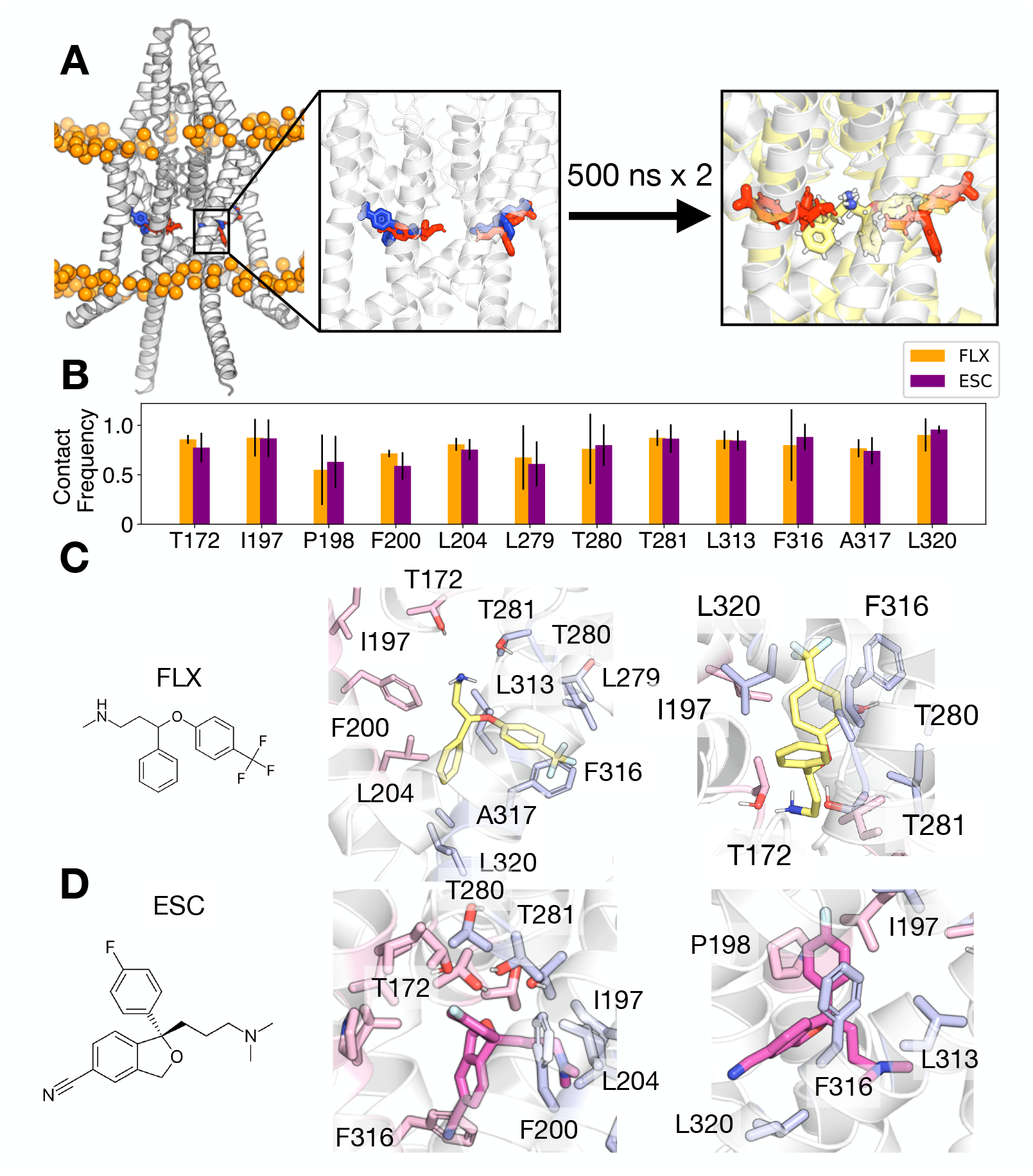
Fluoxetine and escitalopram binding site on the TREK-2 channel. (A) Schematic representation showing the initial set-up of the simulation. The protein is represented in white cartoon and the phosphorus atom of the phospholipid is shown in orange spheres. The crystallographic pose of norfluoxetine is shown in blue, the initial position of fluoxetine is shown in red. After 500 ns of the simulation, the protein and the fluoxetine are shown in pale yellow. (B) Contact probability of residues which are in contact with escitalopram (ESC - purple) and fluoxetine (FLX - orange). Residues with greater than 0.6 contact probability with either fluoxetine, and escitalopram are shown. Error bars show the standard deviation. (C,D) Binding conformation of each fluoxetine (C - yellow) and escitalopram (D - purple) in the binding site. Amino acid residues from each chain are shown in pale blue and pink.

### Fluoxetine and Escitalopram block ion permeation across the pore

We then calculate whether fluoxetine and escitalopram can block the permeation of potassium ions across TREK2. First, we assessed the hydration of the pore cavity in the absence and in the presence of the drugs after 500 ns of the simulation. Here, we showed that the cavity of the TREK2 channel is dehydrated when the drug is present and is fully hydrated without the drug (Figure 2A-C). In the case of escitalopram, the pore cavity is fully hydrated up to the drug molecule. However, in the presence of fluoxetine, the cavity starts to dehydrate from the lower position of the pore, a region which is commonly a bundle crossing gate on the other potassium channel. This observation led to a hypothesis that lipids may block the potassium permeation and dry up the permeation cavity, leading to the energy barrier of potassium permeation (Figure 2B). Previous studies have suggested that lipid fenestration may block ion permeation of the TREK-2 channel, and thus, we calculated whether there is any lipid at the bottom of the pore and whether fluoxetine increases the probability of lipid blocking of the TREK2 channel. We show that palmitoyl oleic phosphatidylcholine (POPC) binds to the bottom of the pore, primarily by making contact with R328 (Figure 2D) and thus, may contribute to the factor of the dehydration of the pore. We then compare this with the simulations with escitalopram or simulations without any drugs. Here, we showed that fluoxetine increases the short-range contact (0.35 nm) that R328 makes with the POPC molecule compared to the simulation without the fluoxetine or with escitalopram in the binding site. Here, we show that in the simulation with fluoxetine, R328 makes more contact with POPC molecules (Figure 2E). Together, this suggested that fluoxetine induces POPC to block the pore and, thus, form parts of the potassium ion blocking mechanism.

**Figure 2.**
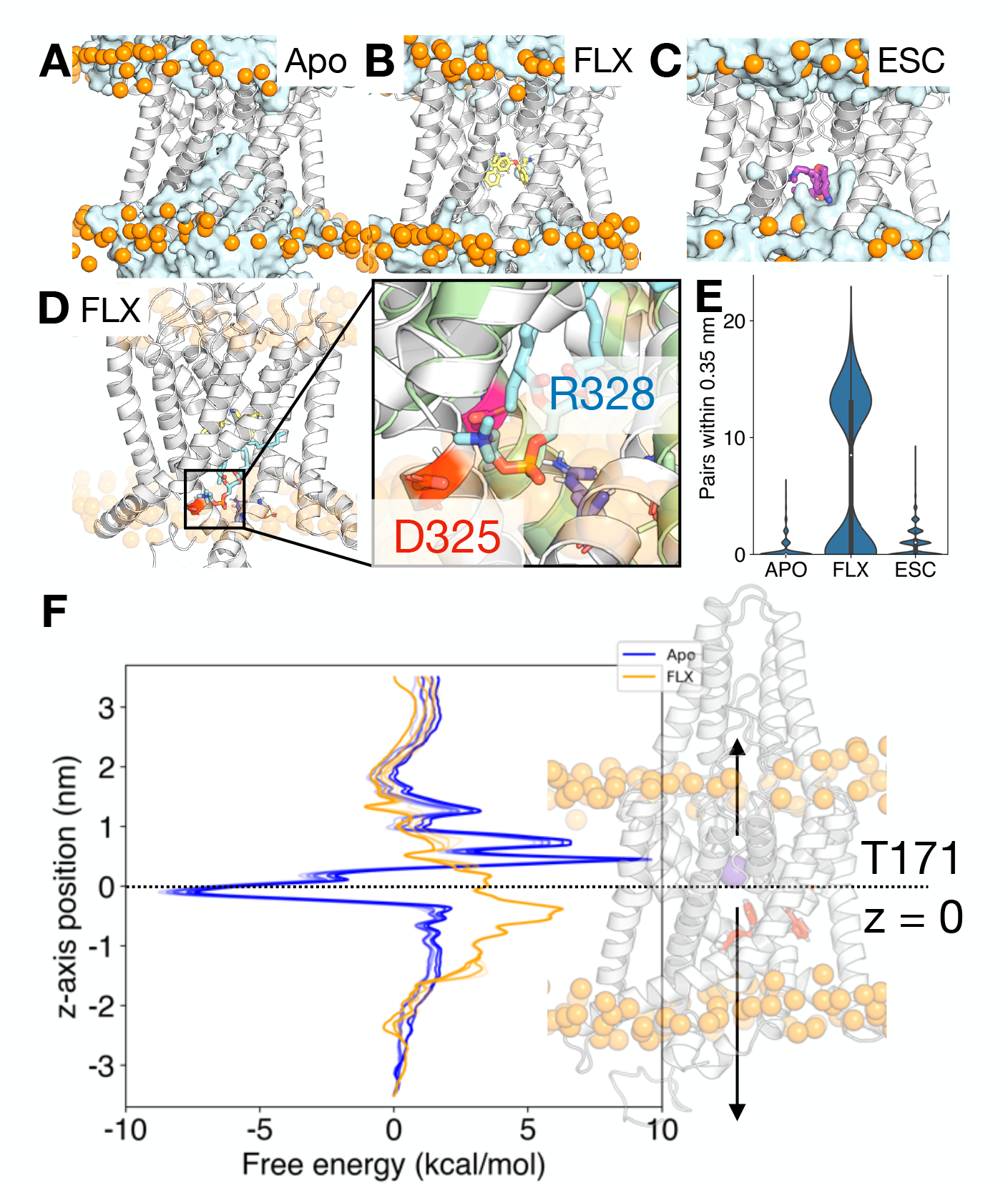
Blocking of the TREK-2 channel by fluoxetine and escitalopram. (A,B,C) Water permeation through the TREK-2 channel in (A) Apo (B) Fluoxetine bound (C) Escitalopram bound. Each panel shows a snapshot at 500 ns from one of the replica. Water is shown as a cyan surface. The TREK-2 channel is shown as white cartoon and the phosphorus atom of the phospholipid is shown in orange spheres. Fluoxetine is shown as yellow sticks and escitalopram as purple sticks. (D) A representative snapshot showing POPC (cyan sticks) binding to R328. (E) Violin plots describing the population of pairwise interaction between POPC and R328 within 0.35 nm in the presence of fluoxetine, escitalopram, or without any of them. (F) Accelerated weighted histogram free energy profile with fluoxetine (orange) and without (blue). The z-axis coordinate is measured with respect to T171 at z = 0

To assess the blockage of potassium ion by the lipid, we conducted accelerated weighted histogram (AWH) enhanced sampling simulations in the presence (FLX) and absence (APO) of fluoxetine (Figure 2E). AWH is an enhanced sampling method, allowing the free energy of the potassium permeation pathway to be studied in the simulation. To do so, we placed one potassium ion at the bottom of the selectivity filter and allowed it to move along the z-axis. Thus, an energy barrier at the fluoxetine binding site would suggest that the molecule can potentially block potassium permeation. Here, we show that the energy barrier started building up from the -2 position on the AWH free energy profile. This position is occupied by PC lipid, and thus, this suggests that PC indeed blocks ion permeation. The barrier height increases as fluoxetine introduces an energy barrier of approximately 5 kcal/mol below the selectivity filter. This energy barrier is absent in the simulation where the drug is not in the binding site. Together, this suggested that fluoxetine induces PC to block the pore and, thus, form parts of the potassium ion blocking mechanism.

In addition to the blockage of potassium ion permeation by lipids and the drugs, we observed three distinct energy barriers in the selectivity filter in the absence of fluoxetine, corresponding to the three potassium positions in the filter. This is a unique feature in the stable potassium ion selectivity filter free energy profile, where the energy from the next potassium ion is required to knock and move the potassium ions forward. However, this distinct feature is absent when the fluoxetine is placed in the binding site. This suggested that fluoxetine may distort the selectivity filter and reduce the selectivity of the TREK2 channel as it blocks the pore.

### Fluoxetine and escitalopram destabilises the selectivity filter of the TREK2 channel

Previous experimental and computational studies have highlighted that the non-conductive state of the K2P channel has a collapsed selectivity filter, which then blocks potassium ion permeation (Schewe et al. 2016; Nematian-Ardestani et al. 2020; Zhang et al. 2022; Rödström et al. 2023). This collapsed selectivity filter is primarily observed when the TREK-2 is in the down or deeper-down conformation. A recent study highlights that fluoxetine inhibits the TREK-2 channel by directly influencing channel gating at the selectivity filter (Proks et al. 2021). Thus, we aim to tackle a similar biological problem using MD simulation. To do so, we removed all potassium ions from the selectivity filter and conducted 500 ns simulations in the absence or the presence of fluoxetine or with escitalopram (n = 4). We then calculated the dihedral angle governing the residues constituting the selectivity filter (T172-G176 - TIGYG and T281-G285 - TVGFG) (Figure 3A, 3B). First, we do not observe any destabilisation of the selectivity filter in the absence of drugs. However, we observed a notable change in the presence of the fluoxetine and escitalopram. On the first pore helix, we do not observe destabilisation at T172. However, in the presence of fluoxetine and escitalopram, we observed destabilisation of the psi angle around I173 and G176 and the dihedral angles around Y175 and G176 (Figure 3A). On the second pore helix, again, the dihedral around T281, V282 and G283 remains relatively similar in the presence and absence of fluoxetine and escitalopram. However, it is striking that fluoxetine and escitalopram distort both dihedral angles around F284 and G285 (Figure 3B). The reason that T172 and T279 make direct contact with the drugs, and thus, allosterically distort the rest of the selectivity conformation. By observing the snapshots of the final frame of the simulation, it is apparent that the simulation with fluoxetine has a more distorted selectivity filter and is, indeed, leading to a slight widening of the selectivity filter (Figure 3C-3E). This observation corresponds to the irregular free energy profile from the AWH calculation (Figure 2A) and thus, may explain the reduction in potassium conduction across the channel in the presence of the fluoxetine.

**Figure 3.**
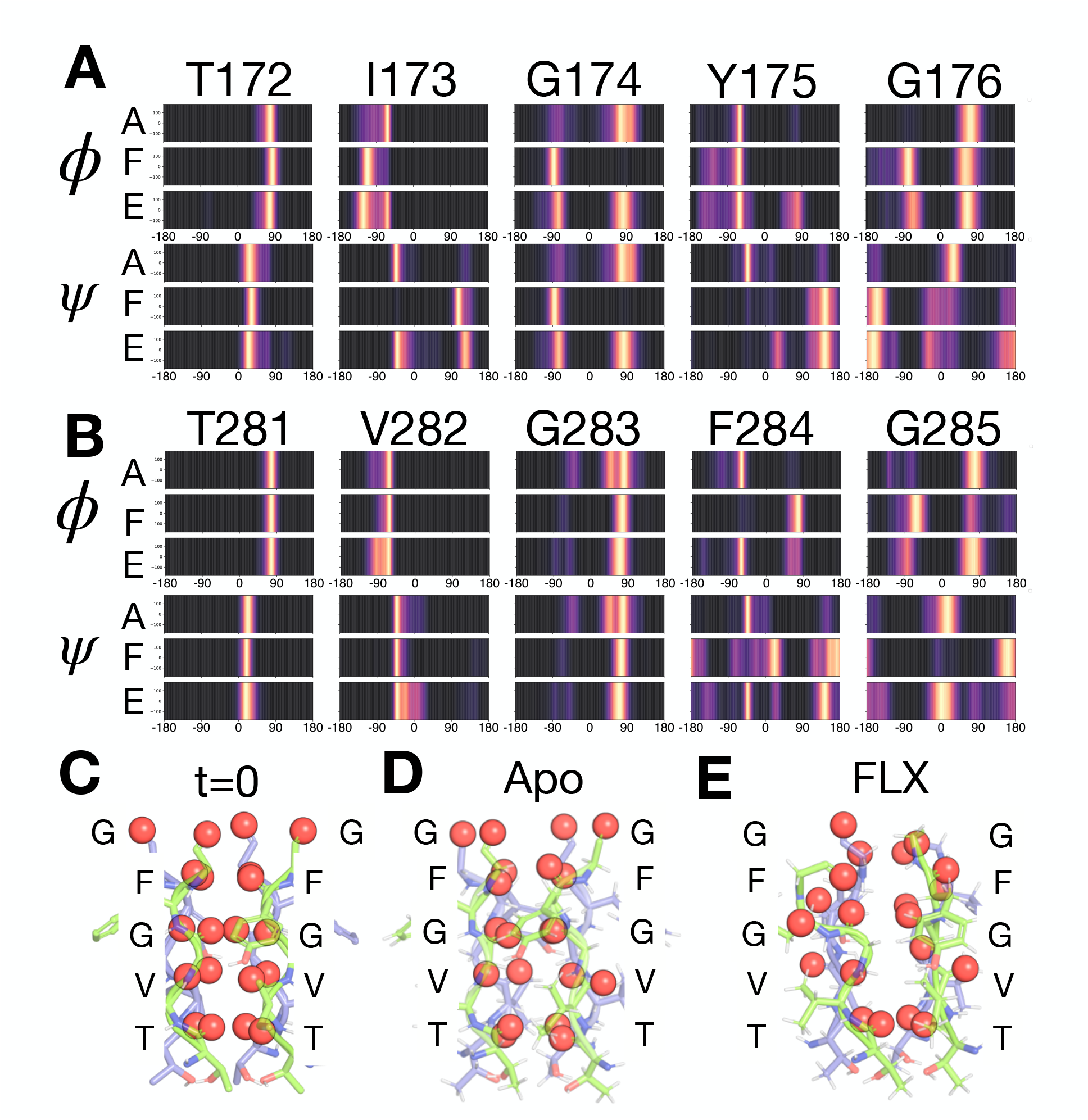
Fluoxetine and escitalopram alter the selectivity filter dynamics of the TREK-2 channel. Distribution of the dihedral angle of the (A) first and the (B) second selectivity filter. Bright spots on the spectrum represent the highly populated region of the specific dihedral angle. (C,D,E) Representative snapshots of the selectivity filter (C) before and after the simulation in the (D) absence of fluoxetine and (E) presence of fluoxetine.

### Fluoxetine and escitalopram stabilise the down-state of the TREK-2 channel

Multiple simulation studies have shown that fluoxetine prevents the conformational transition from the down-conformation to the up-conformation when negative pressure is applied to the system (Aryal et al. 2017; Clausen et al. 2017). This conformational transition can be characterised by three sets of distance measurements, previously defined as fenestration (P198-G324 between two subunits), expansion (M322-G212 on the same subunit) and zipper (W326-R237 of the same subunit) (Figure 4A) (Aryal et al. 2017; Clausen et al. 2017; Zhang et al. 2022). In previous studies, fluoxetine has been shown to stabilise the down state, as the drug makes TREK-2 resistant to the change to the pressure applied to the system (Aryal et al. 2017). Here, we asked whether escitalopram shifted any of the three distances in both the absence of the presence of the pressure and, thus, has any additional role apart from being a traditional pore blocker. Here, we show that by having both fluoxetine and escitalopram in the binding pocket, the three distances remain unchanged. Our work also indicates that fluoxetine and escitalopram do not trigger the transition to the deeper-down state (Figure 4B). When the pressure is applied, all three distances change when the drugs are not in the binding site, which agrees with the previous study (Aryal et al. 2017). However, when either fluoxetine or escitalopram are placed in the binding pocket, the distances remain unchanged (Figure 4C). This suggested that both escitalopram and fluoxetine stabilise the down-state conformation when the pressure is applied, and thus, using a mechanism similar to that of inhibiting the channel.

**Figure 4.**
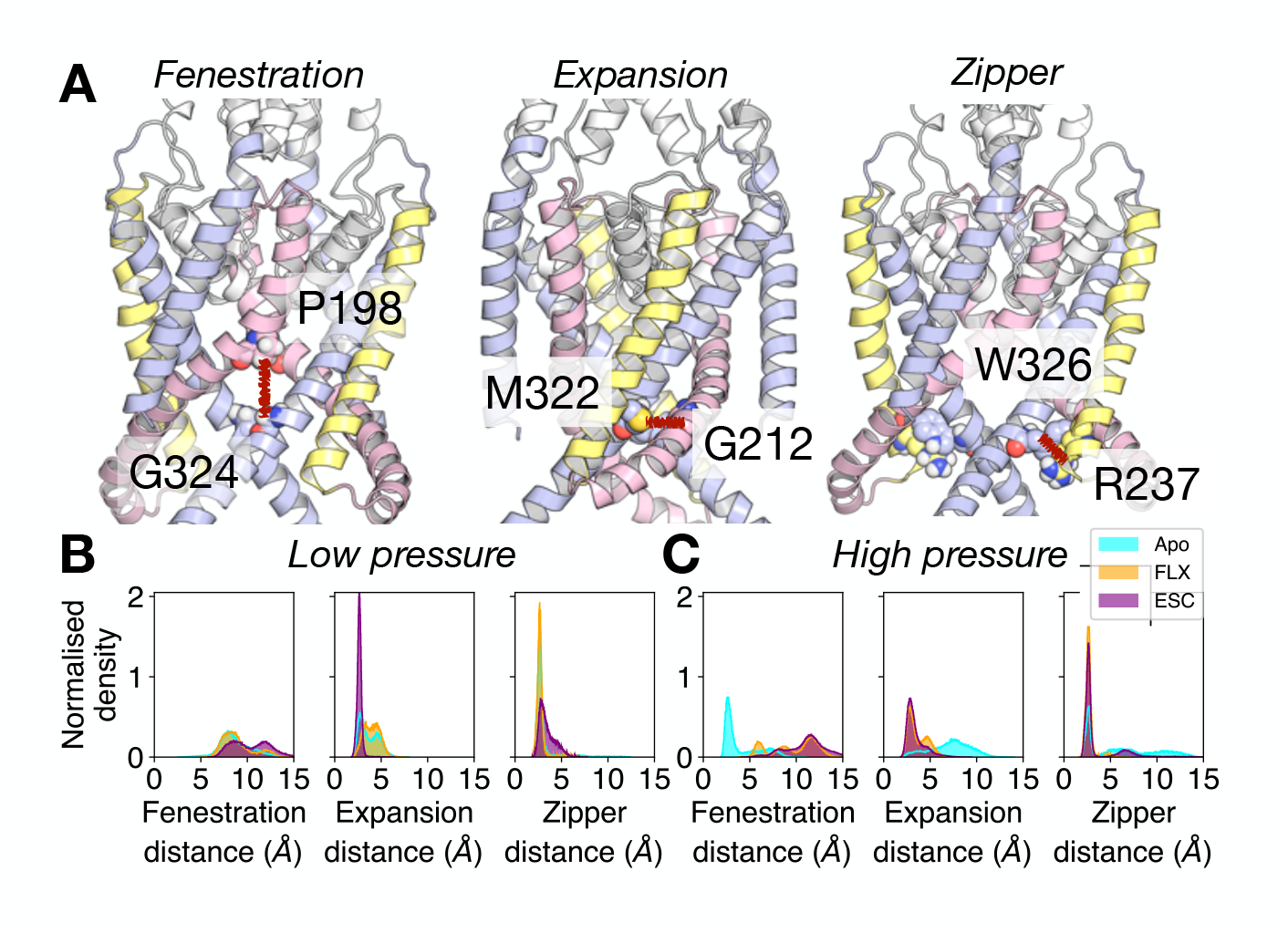
Fluoxetine and escitalopram prevent conformational transition in a stretched bilayer. (A) Cartoon representation of three distances describing down-to-up conformational change, describing (left) fenestration distance or P198-G324 of the opposite subunit, (middle) expansion distance or G212-M312 on the same subunit and (right) zipper distance or W326-R237 on the same subunit. Each colour highlights one set of M1-pore-M2 helix topology. (B,C) Histograms describing the population of the fenestration, zipper and expansion distances with apo (cyan), fluoxetine (orange) or escitalopram bound in the binding site at 1 bar or -50 bar (C).

## Discussion

In this study, we used molecular dynamic simulations to explore the mechanisms of fluoxetine and escitalopram as inhibitors of the TREK-2 channel. By placing both drugs in the fluoxetine binding pocket, we show that fluoxetine and escitalopram share a binding pocket at the lipid fenestration site of TREK-2 (Dong et al. 2015). This site is highly conserved between both TREK-1, TREK-2 and other K2P channels (Schmidpeter et al. 2023). Similar to the previous studies on TWIK-1 channel, our simulation shows that either fluoxetine or escitalopram binding to this site prevents the transition from the down conformation to the up conformation when the pressure is applied to the system (Figure 5) (Aryal et al. 2017). Our simulation also shows that both drugs translocate to the pore region and block the pore. Electrophysiological experiments have demonstrated that escitalopram blocks the TREK2 channel in a voltage-independent manner (Park et al. 2016). Mutations on the escitalopram binding site (T142A, T251A, L174A and L289A) on the TREK1 channel reduce escitalopram inhibition and validate the observed binding site in the simulation (Al-Moubarak, Veale, and Mathie 2013). Our simulation also shows that escitalopram blocks the pore hydration and does not increase the distortion of the selectivity filter. This suggested that escitalopram behaves as a traditional pore blocker when the TREK2 channel is in the down conformation.

**Figure 5.**
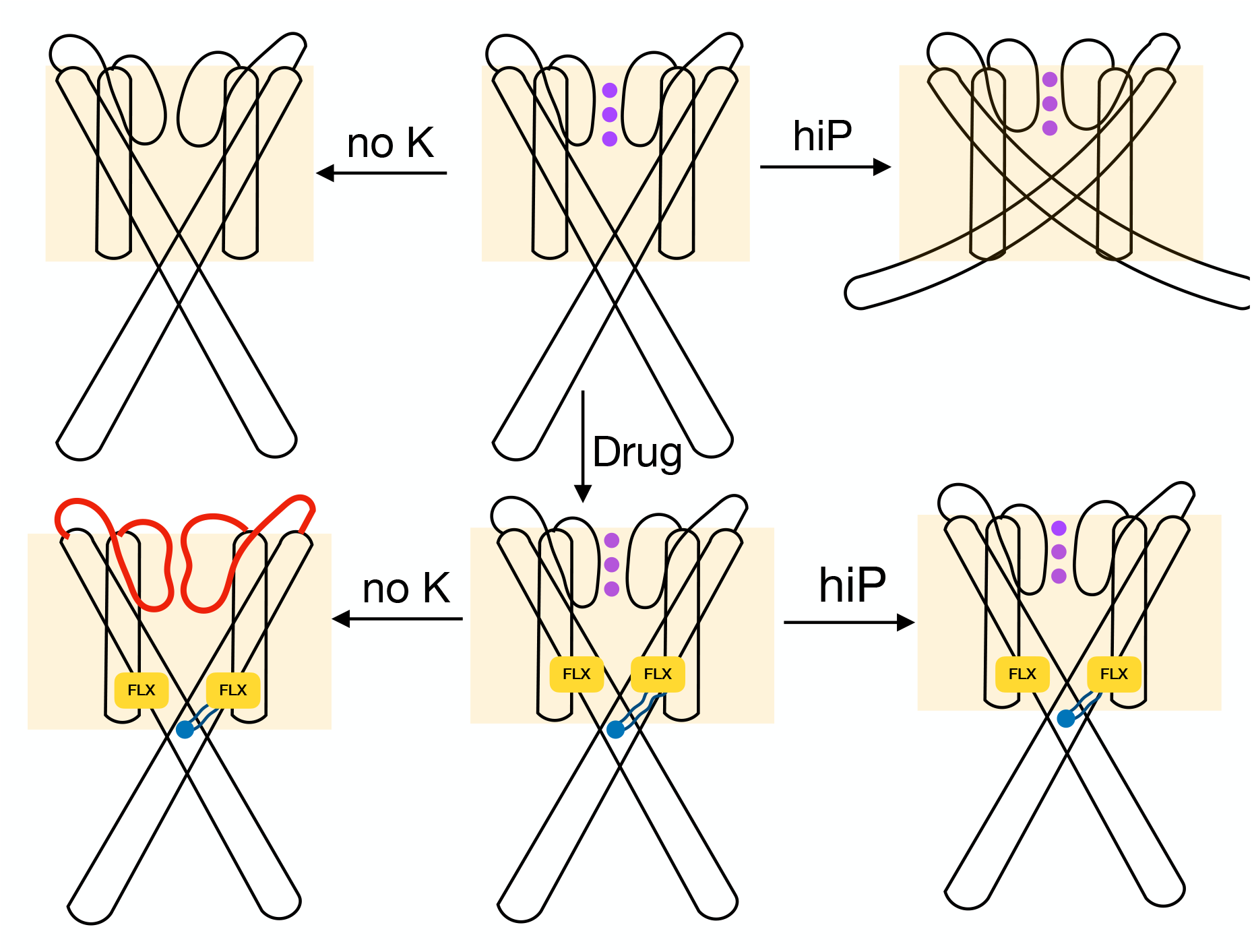
Schematic representation of fluoxetine and escitalopram mechanism. Schematic representation showing how pressure or SSRIs interact with the TREK-2 channel. The phospholipid bilayer is shown in orange. Fluoxetine is shown as yellow rectangles and escitalopram as pink rectangles. Potassium ions in the selectivity filter are shown as purple circles. The distorted selectivity filter is shown in red. POPC is shown in blue.

Single-channel recording studies of the TREK2 channel highlighted that norfluoxetine is not a pore blocker but inhibits the channel through conformational stabilisation and selectivity filter distortion (Proks et al. 2021). Structural work proposes that one of the possible inhibitory mechanisms of fluoxetine is through lipid fenestration site (Lolicato et al. 2017a). This lipid fenestration site and the migration of lipids to the pore is similar to what we have previously observed in coarse-grained molecular dynamics simulation with phosphatidic acid (Panasawatwong, Pipatpolkai, and Tucker 2022). This site is nearby to the site where dodecylmaltoside (DDM) detergent, and phosphatidylethanolamine (PE) were observed in the very recent down conformation structure of TREK-1 (Schmidpeter et al. 2023). Our simulation and free energy profile suggest that blocking of the channel by fluoxetine involves PC interaction with R328, leading to pore dehydration. This is supported by a voltage-independent blocking of the TREK-2 channel by fluoxetine and norfluoxetine (Kennard et al. 2005; Proks et al. 2021). We also observed the distortion of the selectivity filter of the TREK2 channel only in the presence of fluoxetine, which supports the single-channel recording experiment (Proks et al. 2021). We postulate that this will also be the same for escitalopram. Together, our work proposes that the fluoxetine mechanism of TREK2 inhibition involves the distortion of the selectivity filter observed from the simulation as the rate-determining step rather than the channel’s direct pore blocking and lipid blocking. Understanding these mechanisms could be extended to other K2P channel families, thus leading to a greater understanding of the treatment of depression.

## Materials and Methods

### Model building and simulation set-up

The TREK-2 down conformation sequence residue 73-352 was taken from UniProt: P57789-4. The extended helical structure at the C-terminus (residue 331-352) is modelled based on the TREK-1 up conformation structure (PDB ID: 6CQ6) (Lolicato et al. 2017a) by superimposing the helical structure onto the model, whereas the rest of the structure TREK-2 channel is modelled based on the down conformation with norfluoxetine bound TREK-2 (PDB entry: 4xdk) (Dong et al. 2015) using SwissModel. Both fluoxetine and citalopram are placed in the norfluoxetine binding site by superimposing the molecules onto the structure. The structure is then embedded in the 100% POPC bilayer on both leaflet and then energy minimised and equilibrated based on CHARMM-GUI protocol (Wu et al. 2014). The system was then energy minimised and equilibrated using the standard six steps CHARMM-GUI equilibration protocol. This includes the following set-up: The protein backbone was restrained at the force constant of 4000, 2000, 1000, 500, 200 and 50 kJmol^-1^nm^-2^, the protein side chain was restrained the force constant of 2000, 1000, 500, 200, 50, and 0 kJmol^-1^nm^-2^, the lipids non-H atoms were restrained at the force constant of 1000, 400, 400, 200, 40 and 0 kJmol^-1^nm^-2^ and the dihedral restraint was set at the force constant of 1000, 400, 200, 200, 100 and 0 kJmol^-1^rad^-2^. The simulations were equilibrated with a 1 fs timestep for 125 ps for the first three steps, and then to a 2 fs timesteps for 500 ps the next two, and 5 ns for the final step. The first two steps were conducted with the NVT ensemble where the last four were conducted with the NPT ensemble. All equilibration runs were conducted at 310 K using Berendsen thermostat (Berendsen et al. 1984). In all NPT ensemble equilibration, the pressure was maintained at 1 bar using Berendsen barostat (Berendsen et al. 1984). The production runs were conducted for 500 ns under 310 K using the v-rescale thermostat (Bussi, Donadio, and Parrinello 2007). The pressure of all systems was maintained using a c-rescale barostat (Bernetti and Bussi 2020). All simulations with no externally biased set-up and a number of replicates are shown in Table 1. All simulations are conducted with CHARMM36m forcefield. All system contains 1 protein, 211 POPC lipid, 66 potassium ion, 70 chloride ions (0.15 M KCl) and, 24776 TIP3P water molecules in a 9 nm x 9 nm x 13.3 nm box. All simulations were conducted using GROMACS-2022 (Abraham et al. 2015).

**Table 1:** All simulations set up in this study.

| Drugs | Simulation lengths (ns) | Potassium ion in the selectivity filter | Pressure (bar) | Number of replicates |
| --- | --- | --- | --- | --- |
| None (Apo) | 500 | 3 | 1 | 2 |
| None (Apo) | 500 | 3 | -50 | 2 |
| None (Apo) | 500 | 0 | 1 | 4 |
| Fluoxetine (FLX) | 500 | 3 | 1 | 2 |
| Fluoxetine (FLX) | 500 | 3 | -50 | 2 |
| Fluoxetine (FLX) | 500 | 0 | 1 | 4 |
| Escitalopram (ESC) | 500 | 3 | 1 | 4 |
| Escitalopram (ESC) | 500 | 3 | -50 | 4 |
| Escitalopram (ESC) | 500 | 0 | 1 | 4 |

### Accelerated weighted histogram

The free energy of potassium permeation across the TREK-2 channel was calculated using an accelerated weighted histogram (AWH) (Lindahl, Lidmar, and Hess 2014). The TREK-2 channel has no potassium ions in the selectivity filter. The pressure is maintained at 1 bar. We applied an independent AWH bias for each equilibrated structure and simulated 4 walkers for 500 ns (total of 2000 ns), sharing bias data and the same target distribution. The bias acts on the z-axis defined using the centre-of-mass of the K ion and the T171 at the bottom of the selectivity filter of the TREK-2 channel, aiming for a flat target distribution. The sampling interval was 3.5 nm above and below the T171. The system initialised with the average free energy error of 20 kJ/mol, with the diffusion coefficient at 0.0002 nm^2^/ps at a force constant of 12800 kJ mol^−1^ nm^−2^. The harmonic potential was applied on all C-alpha toms at 1000 kJ mol^−1^ nm^−2^. The convergence was assessed by gaining a relatively similar free energy landscape over the last 50 ns of the simulations and a flat-target distribution.

## Acknowledgements

We thank Assoc. Prof. Lucie Delemotte and her laboratory for their support and advice throughout the project. We also thank Prof. Stephen Tucker and Asst. Prof. Puey Ounjai for their critical reading of the manuscript. The MD simulations were performed on resources provided by the Swedish National Infrastructure for Computing (SNIC) on Dardel at the PDC Center for High Performance Computing (PDC-HPC), and ASPIRE2a HPC.

## Author contributions

T.P. and N.P. designed, performed research and analysed the data. T.P. wrote and revised the manuscript.

